# Comparison of Commercially Available Thermostable DNA Polymerases with Reverse-Transcriptase Activity in Coupled Reverse-Transcription Polymerase Chain Reaction Assays

**DOI:** 10.1101/2024.08.03.606463

**Authors:** Evgeniya V. Smirnova, Konstantin A. Blagodatskikh, Ekaterina V. Barsova, Dmitriy A. Varlamov, Vladimir M. Kramarov, Konstantin B. Ignatov

**Author notes:** **Correspondence:** Konstantin B. Ignatov, Vavilov Institute of General Genetics, Russian Academy of Sciences, Moscow, Russia.

## Abstract

Reverse-transcription polymerase chain reaction (RT-PCR) is an important tool for the detection of target RNA molecules and the assay of RNA pathogens. Coupled RT-PCR is performed with an enzyme mixture containing a reverse transcriptase and a thermostable DNA polymerase. To date, several biotechnological companies offer artificial thermostable DNA polymerases with a built-in reverse-transcriptase activity for use in the coupled RT-PCR instead of the enzyme mixtures. Here, we compared the artificial DNA polymerases and conventional enzyme mixtures for the RT-PCR by performing end-point and real-time RT-PCR assays using severe acute respiratory syndrome-related coronavirus 2 (SARS-CoV2) RNA and endogenous mRNA molecules as templates. We found that the artificial enzymes were suitable for different RT-PCR applications, including SARS-CoV2 RNA detection, but not for long-fragment RT-PCR amplification.

## 1 Introduction

The coronavirus disease 2019 (COVID-19) pandemic demonstrated the importance of robust and relatively inexpensive methods, including reverse-transcription polymerase chain reaction (RT-PCR), for the detection of RNA viruses. RT-PCR and reverse-transcription quantitative polymerase chain reaction (RT-qPCR) are the most important methods for the detection and quantification of target RNA sequences that are widely used for monitoring expression levels of mRNA and RNA biomarkers and the assay of RNA pathogens such as RNA viruses. Traditionally, RT-PCR is performed with two enzymes: a reverse transcriptase (e.g., MMLV or AMV revertase) for cDNA synthesis and a thermostable DNA-dependent DNA polymerase (e.g., Taq or Tth polymerase) for the PCR amplification of cDNA. However, using two separate enzymes complicates a reaction system and increases the cost of RT-PCR. Additionally, reverse transcriptases inhibit Taq polymerase activity and decrease the efficiency of PCR amplification (Sellner et al., 1992; Fehlmann et al., 1993; Chumakov, 1994; Suslov and Steindler, 2005). These problems could be solved by using a single enzyme in a coupled RT and PCR amplification. Unfortunately, natural reverse transcriptases are not thermostable and cannot be used for PCR. On the contrary, Taq polymerase has very low RT activity and cannot be used for cDNA synthesis under normal conditions. The first attempts to tackle these problems were made in the 1990s when Mn^2+^ was used instead of Mg^2+^ to stimulate RT activity of Taq and Tth DNA polymerases (Tse and Forget, 1990; Myers and Gelfand, 1991; Grebennikova et al., 1995; Grabko et al., 1996). Unfortunately, this improvement required routine optimization of the reaction buffer and did not provide high efficiency of the RT step of the reaction. Another approach to create a single enzyme that could be used in RT and PCR was based on the protein engineering of the thermostable Family A (PolA) DNA polymerases, mainly Taq DNA polymerase. Several research groups have developed a selection of enzymes using different modifications of thermostable DNA polymerases, which are suitable for performing RT-PCR in a single enzyme mode.

The first articles describing modifications of Taq polymerase for enhancing reverse-transcriptase activity were published in 2006 (Ong et al., 2006; Vichier-Guerre et al., 2006; Sauter and Marx, 2006). The most suitable enzymes for RT-PCR applications were mutants of Taq polymerase, which were described in 2010 by Prof. Andreas Marx and his research group from University of Konstanz, Germany (Kranaster et al., 2010; Blatter et al., 2013). The optimized mutant of Taq polymerase contained four substitutions in the polymerase domain: S515R, L459M, I638F, and M747K. This development was commercialized by myPOLS Biotec GmbH (Germany) as the RevTaq DNA polymerase and Volcano master mix. The created polymerase was successfully applied for the detection of SARS-CoV2 RNA (Kuiper et al., 2020; Babler et al., 2023).

The scientists affiliated with QIAGEN (Beverly MA, USA) created Magma DNA polymerase with an improved reverse-transcriptase activity (Heller et al., 2019). Magma polymerase is a fusion enzyme between the viral 3173 PyroPhage DNA polymerase and the 5`→3` nuclease domain of Taq DNA polymerase. PyroPhage DNA polymerase is a thermostable viral Family A polymerase with inherent reverse-transcriptase activity. The conjugation of the PyroPhage polymerase with the 5`→3` nuclease domain of Taq DNA polymerase confers compatibility with TaqMan probes. Experiments demonstrated that Magma could be used to carry out RT-PCR (Heller et al., 2019); however, this enzyme has not been commercialized.

Another modification of Taq polymerase was described in 2021 by Prof. Wayne Barnes and his colleagues from DNA Polymerase Technology, Inc. (MO, USA). They found that a single substitution D732N in the polymerase domain of Taq polymerase conferred strand-displacement and reverse-transcriptase activities on the enzyme (Barnes et al., 2021). The presence of strand-displacement activity allows the enzyme to overcome the secondary structure of RNA template; all natural reverse transcriptases possess a strong strand-displacement activity (Whiting et al., 1994).

The D732N mutant of Taq polymerase was commercialized by DNA Polymerase Technology, Inc. as the OmniTaq2 DNA polymerase.

A new variant of Tth DNA polymerase with reverse-transcriptase activity was created by introducing two amino acid substitutions I709K and I640F into the enzyme (Luo et al., 2023).

The ReverHotTaq DNA polymerase, offered by Bioron GmbH (Germany), is another thermostable enzyme with strand-displacement and reverse-transcriptase activities. According to the description of the ReverHotTaq, it was obtained by incorporation of the fragments from the Bst DNA polymerase amino acid sequence into the active site of Taq DNA polymerase. The Bst polymerase has both strand-displacement and a reverse-transcriptase activities but is not thermostable enough for performing PCR (Ignatov et al., 2014). Thus, the engineered ReverHotTaq combines the advantages of both parent enzymes: strand-displacement and reverse-transcriptase activities from the Bst polymerase as well as the 5’→3’ DNA polymerase and 5’→3’ exonuclease activities with high thermostability from the Taq polymerase.

The use of a single enzyme for coupled RT-PCR could greatly simplify the reaction system and decrease the cost of RT-PCR tests. Unfortunately, only a few examples of practical application of the modified thermostable polymerases for RT-PCR tests are currently known in the field (Kuiper et al., 2020; Babler et al., 2023) and a direct comparison of the enzymes *inter se* has not yet been done.

In the present work, we compare three different commercially available thermostable DNA polymerases with reverse-transcriptase activity (RevTaq, OmniTaq2, and ReverHotTaq) by performing end-point and real-time RT-PCR assays of SARS-CoV2 RNA and mRNA molecules. Conventional M-MLV/Taq polymerase mixtures for OneTube RT-PCR were used as standards.

## 2 Materials and Methods

### 2.1 Enzymes and reagents

The following materials were obtained from these sources: 100X OmniTaq2 DNA polymerase and 10X Taq Mutant Reaction Buffer from DNA Polymerase Technology, Inc. (St. Louis, MO, USA); 50X RevTaq DNA polymerase and 5X Volcano Reaction Buffer from myPOLS Biotec GmbH (Konstanz, Germany); 50X ReverHotTaq polymerase and 5X Reaction Buffer from Bioron GmbH (Römerberg, Germany); OneTaq One-Step RT-PCR Kit from New England BioLabs, Inc. (Ipswich, MA, USA); 5X OneTube RT-PCR-Mix from Evrogen (Moscow, Russia); and dNTPs from Bioline Limited (London, UK). Oligonucleotide primers and TaqMan probes were synthesized by Syntol JSC (Moscow, Russia). SARS-CoV-2 viral RNA isolated from nasopharyngeal swabs was provided by Syntol and verified by using the SARS-CoV-2 RT-qPCR Detection Kit (Syntol JSC, Moscow). The real-time amplification reactions were performed using a CFX96 Touch Real-Time Detection System (Bio-Rad Laboratories, Inc., Hercules, CA, USA). EvaGreen intercalating dye was supplied by Syntol JSC (Moscow, Russia).

Human total RNA was isolated from human embryonic kidney 293 (HEK 293) cells by using TRIzol reagent (ThermoFisher Scientific, Inc., Carlsbad, CA, USA) and then subjected to treatment with RNase-Free DNase I (RNase-Free DNase Set, Qiagen GmbH, Hilden, Germany), followed by column purification (RNeasy Mini Kit, Qiagen GmbH, Hilden, Germany). Total RNA concentration was measured with a Qubit RNA High Sensitivity kit and Qubit-4 fluorimeter (ThermoFisher Scientific, Inc., Carlsbad, CA, USA).

### 2.2 Coupled end-point RT-PCR

End-point RT-PCR assays were carried out with 10 ng, 1 ng, 100 pg, or 10 pg of human total RNA or without RNA as a negative control. The reaction mixtures (25 *μ*L) contained 0.25 mM dNTP (each), two primers (0.3 *μ*M, each), 1X concentration of RevTaq (0.5 *μ*L) or OmniTaq2 (0.25 *μ*L) or ReverHotTaq (0.5 *μ*L) or One Taq One-Step Enzyme Mix (1 *μ*L), and 1X concentration of the appropriate reaction buffer. The following primers were used for the amplification of beta-2-microglobulin cDNA fragments: *B2M-f1* (5’-CTGCCGTGTGAACCATGTGA) and *B2M-r1* (5’-CAATCCAAATGCGGCATCTTC) for the 105 bp amplicon; *B2M-f2* (5’-TGTAAGCAGCATCATGGAGGTT) and *B2M-r2* (5’-TGCTCAGATACATCAAACATGG) for the 203 bp amplicon; *B2M-f1* and *B2M-r3* (5’-CTCTGCTCCCCACCTCTAAGTT) for the 317 bp amplicon.

The following conditions were used for end-point RT-PCR with RevTaq, OmniTaq 2, and ReverHotTaq DNA polymerases: (a) initial denaturation at 75 °C for 2 min.; (b) reverse transcription at 60 °C for 10 min., then 68 °C for 30 min.; (c) denaturation at 92 °C for 1 min.; (d) PCR amplification by 38 thermocycles at 92 °C for 30 s, 60 °C for 30 s, and 68 °C for 30 s. For RT-PCR with the OneTaq One-Step RT-PCR enzyme mixture, the following conditions were applied: (a) RT at 48 °C for 30 min.; (b) denaturation at 92 °C for 1 min.; (c) PCR amplification by 38 thermocycles at 92 °C for 30 s, 60 °C for 30 s, and 68 °C for 30 s.

### 2.3 Coupled real-time RT-qPCR with an intercalating dye

Real-time RT-qPCR assays with EvaGreen DNA-binding dye were carried out with 1 ng, 100 pg, or 10 pg of human total RNA or without RNA as a negative control. Polymerases RevTaq, OmniTaq 2, ReverHotTaq, and OneTube RT-PCRMix were used for the assays. The reaction mixtures were identical to the mixtures of end-point RT-PCR described above, except the primers for target amplification. Additionally, the reaction mixtures contained 0.4X EvaGreen DNA-binding dye. Coupled reverse transcriptions and real-time amplifications of a 109 bp fragment of human *GAPDH* cDNA were performed with primers *GAPDH_d* (5’-GTGGTCTCCTCTGACTTCAACAG) and *GAPDH_r* (5’-CGTTGTCATACCAGGAAATGAGCTTG).

For RT-qPCR with RevTaq, OmniTaq 2, and ReverHotTaq polymerases, we used the following conditions: (a) initial denaturation at 75 °C for 2 min.; (b) reverse transcription at 62 °C for 10 min., then 68 °C for 15 min.; (c) denaturation at 92 °C for 2 min.; (d) PCR amplification by 40 thermocycles: 92 °C for 15 s, 60 °C for 15 s, and 68 °C for 15 s. For RT-qPCR with OneTube RT-PCRmix, we applied (a) reverse transcription at 50 °C for 20 min.; (b) denaturation at 92 °C for 2 min.; (c) PCR amplification by 40 thermocycles: 92 °C for 15 s, 60 °C for 15 s, and 68 °C for 15 s.

### 2.4 Coupled real-time RT-qPCR with a TaqMan probe

TaqMan probe-based RT-qPCRs targeting SARS-CoV-2 viral RNA were performed using OneTube RT-PCRMix and polymerases: RevTaq, OmniTaq 2, and ReverHotTaq. The reactions were carried out over a 5-log range of input template RNA concentrations. The RNA isolated from nasopharyngeal swabs containing SARS-CoV-2 viral genomic RNA (gRNA) was diluted 10, 50, and 250 times and used as a template. The reaction mixtures (25 *μ*L) contained 10 *μ*L of the template RNA, 0.25 mM dNTP (each), two primers (0.4 *μ*M, each), TaqMan probe (0.2 *μ*M), 1X concentration of RevTaq (0.5 *μ*L) or OmniTaq2 (0.25 *μ*L) or ReverHotTaq (0.5 *μ*L) or OneTube RT-PCRMix (5 *μ*L), and 1X concentration of the appropriate reaction buffer. The following primers and probe were used for the RT-qPCR reactions: *CoV-d* (5’-ACCTTAAATTCCCTCGAGGACAA), *CoV-r* (5’-TAGGTAGTAGAAATACCATCTTGGACTGAGA), and *CoV-probe* (5’-(ROX)-TCCAATTAACACCAATAGCAGTCCAGA-(BHQ2)).

The assays with RevTaq, OmniTaq 2, and ReverHotTaq polymerases were performed as follows: (a) initial denaturation at 75 °C for 2 min.; (b) reverse transcription at 62 °C for 10 min., then 68 °C for 15 min.; (c) denaturation at 92 °C for 2 min.; (d) PCR amplification by 50 thermocycles: 92 °C for 15 s, 58 °C for 15 s, and 70 °C for 15 s. The conditions for RT-qPCR with OneTube RT-PCRmix are as follows: (a) reverse transcription at 50 °C for 20 min.; (b) denaturation at 92 °C for 2 min.; (c) PCR amplification by 50 thermocycles: 92 °C for 15 s, 58 °C for 15 s, and 70 °C for 15 s.

### 2.5 Data analysis

All calculations were performed using R software (R Core Team, 2021). Core R language and several extension packages were used, including RDML for raw amplification data manipulation (Rödiger et al., 2017), chipPCR for Cq calculation (Rödiger et al., 2015), and ggplot2 for the generation of plots (Wickham, 2016).

## 3 Results

### 3.1 End-point RT-PCR

The performance of three thermostable DNA polymerases with reverse-transcriptase activity (RevTaq, OmniTaq2, and ReverHotTaq) was tested in end-point RT-PCR assays. OneTaq One-Step Enzyme Mix (NEB), a specially designed enzyme mixture for end-point RT-PCR, was used as a standard. Target sequences of human beta-2-microglobulin (*B2M*) mRNA of different sizes (105 bp, 203 bp and 317 bp) were amplified as described in *Materials and Methods*. For the experiments, we used total human RNA and primers for *B2M* mRNA. As the human genome does not contain any pseudogenes of *B2M*, it is widely used as the endogenous reference gene for the relative quantification of genes of interest (Balaji, S. et al., 2020).

Figure 1 shows that all the enzymes demonstrated similar sensitivity and product yields when amplifying the shortest 105 bp fragment. The efficiency of the RevTaq and ReverHotTaq polymerases was equal to that of the OneTaq One-Step RT-PCR Kit when amplifying the 203 bp fragment. However, the results for the OmniTaq2 polymerase showed that once the length of the cDNA fragments increased from a small 105 bp fragment to a larger 203 bp or 317 bp fragment, the yield of the specific products decreased. When amplifying the largest 317 bp fragment, a tenfold decrease in sensitivity was demonstrated by RevTaq, OmniTaq2, and ReverHotTaq polymerases compared to the OneTaq One-Step RT-PCR Kit. Figure 1 also shows that in all reactions, RevTaq polymerase generates more visible amounts of non-specific products, compared to other enzymes.

**Figure 1.**
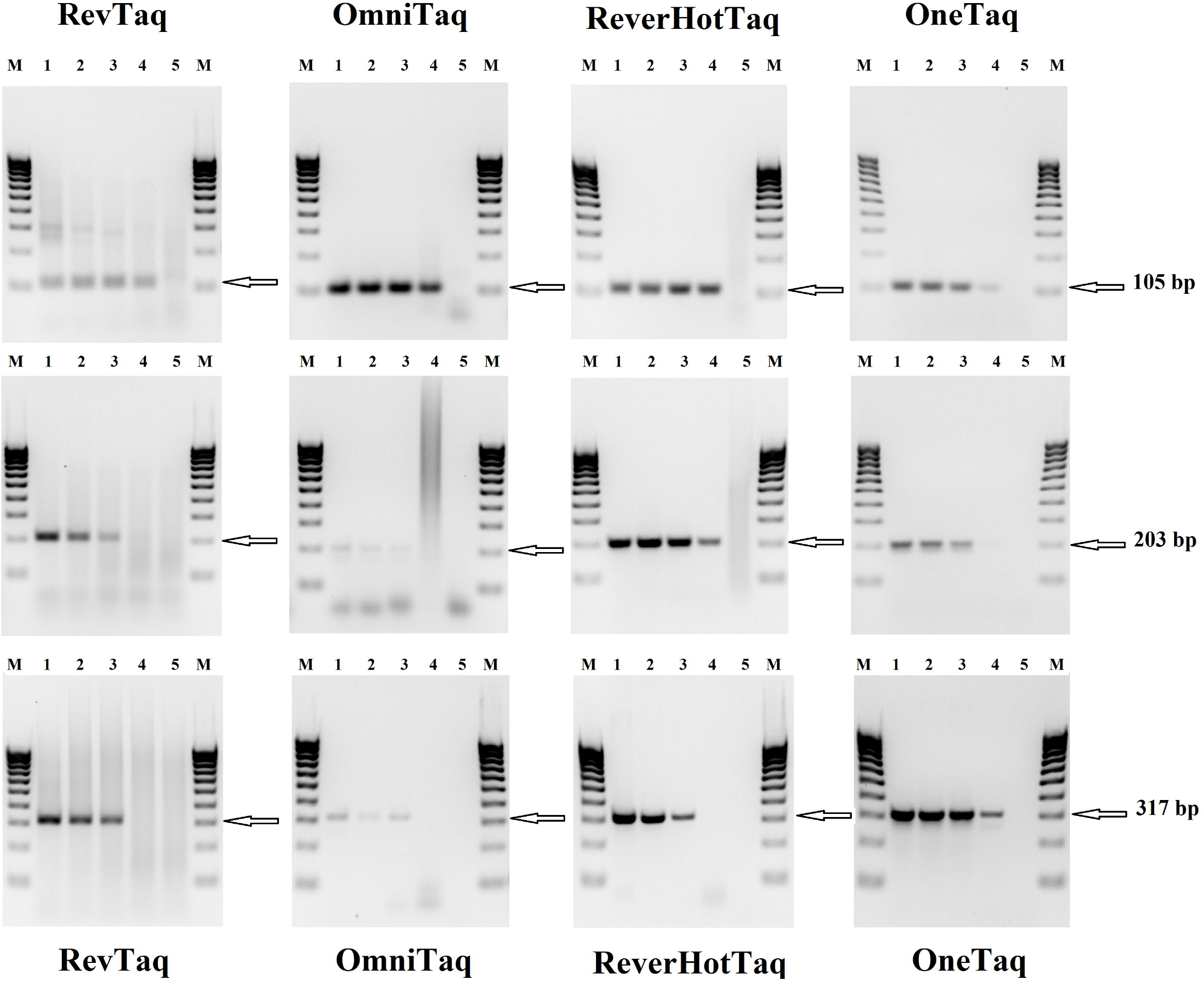
End-point RT-PCR assays with RevTaq, OmniTaq 2, and ReverHotTaq DNA polymerases and OneTaq One-Step Enzyme Mix. The indicated DNA polymerases and OneTaq enzyme mixture were used for the RT-PCR amplification of 105 bp, 203 bp, and 317 bp fragments (indicated by arrows) of the human beta-2-microglobulin mRNA sequence. The reaction mixtures contained a human total RNA as a template: 10 ng – lane 1; 1 ng – lane 2; 100 pg – lane 3; 10 pg – lane 4; no template control (NTC) – lane 5.

### 3.2 Real-time RT-qPCR with an intercalating dye

The performance of RevTaq, OmniTaq2, and ReverHotTaq DNA polymerases was compared to that of the OneTube RT-PCRMix in real-time RT-qPCR with EvaGreen dye. The OneTube RT-PCRMix containing a reverse transcriptase and Taq polymerase is a specially designed enzyme mixture for coupled real-time RT-qPCR. The target 109 bp sequence of human *GAPDH* mRNA was amplified from the series of dilutions of human total RNA as described in *Materials and Methods. GAPDH* is the standard endogenous reference gene for expression studies (Balaji, S. et al., 2020). Figure 2 and Table 1 show the results of the RT-qPCRs; ReverHotTaq DNA polymerase and OneTube RT-PCRMix had similar efficiency with different template concentrations. RevTaq polymerase showed similar results to ReverHotTaq when 1 ng and 100 pg of the total RNA were used but generated non-specific products at low template concentration (Fig. 2B, melting curves), which prevented the detection of *GAPDH* mRNA in 10 pg of human total RNA (Fig. 2B). OmniTaq2 DNA polymerase allowed the RT-qPCR assay to be carried out with all the RNA template concentrations (Fig. 2C) but demonstrated significantly lower efficiency of amplification than the other enzymes, which was seen as a 3–6 cycle lag in the RT-qPCR (Fig. 2, Table 1). In the no template control (NTC) reactions, all enzymes generated signals from non-specific products (Fig. 2).

**Table 1.**
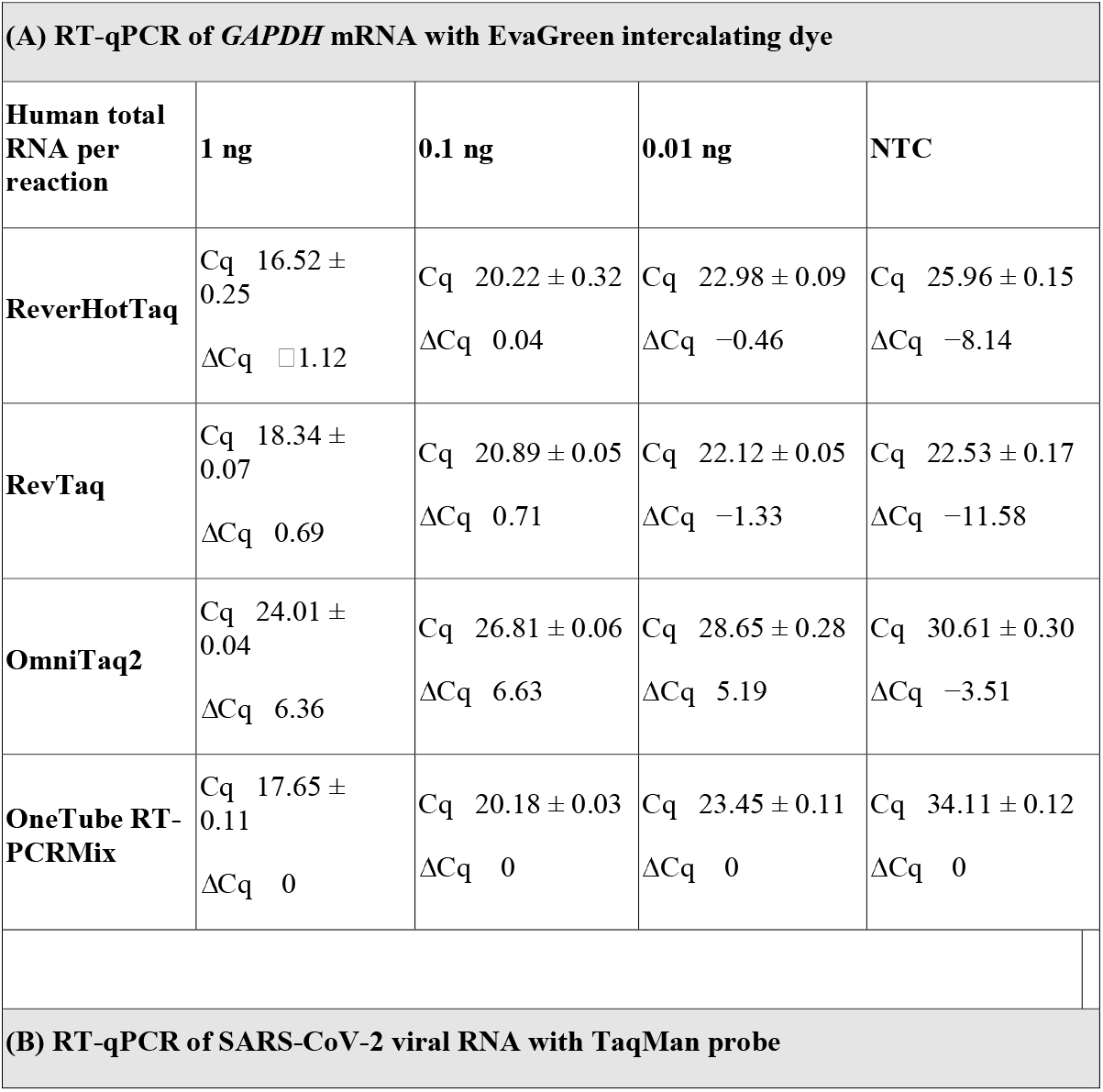
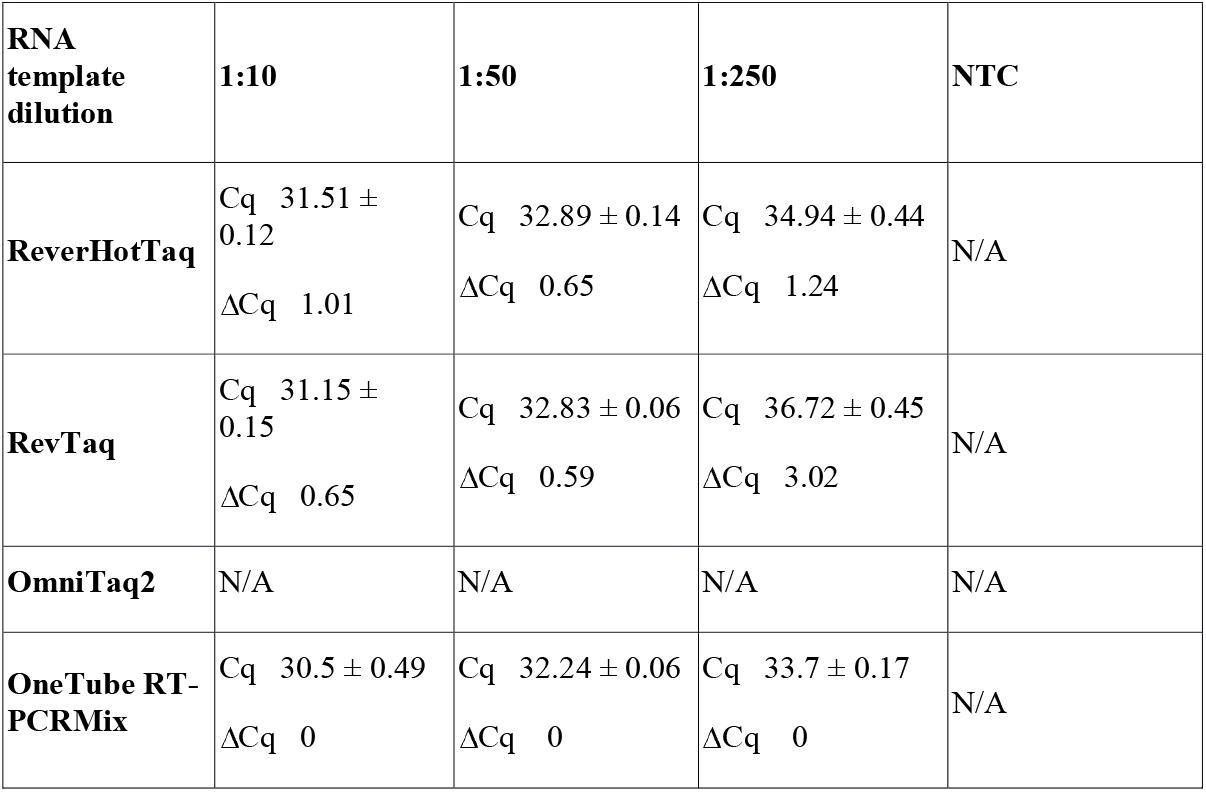
Comparison of the indicated enzymes in RT-qPCR with EvaGreen intercalating dye (A) and with TaqMan probe (B). The mean quantification cycles (Cq) ± standard deviation of three replicates for the indicated template total RNA amounts (NTC–no template control) are provided. ΔCq is the difference between the Cq of the RT-qPCR with the indicated polymerase and control OneTube RT-PCR-Mix for each template dilution. (A) The real-time amplifications of *GAPDH* mRNA fragment with EvaGreen intercalating dye were carried out using 10-fold dilutions of the human total RNA from 1 ng to 0.01 ng as a template, or no template control (NTC). (B) The TaqMan prob RT-qPCR assays of SARS-CoV-2 viral RNA were carried out with 10, 50, and 250 times diluted RNA isolated from SARS-CoV-2-positive nasopharyngeal swabs as a template or no template control (NTC).

**Figure 2.**
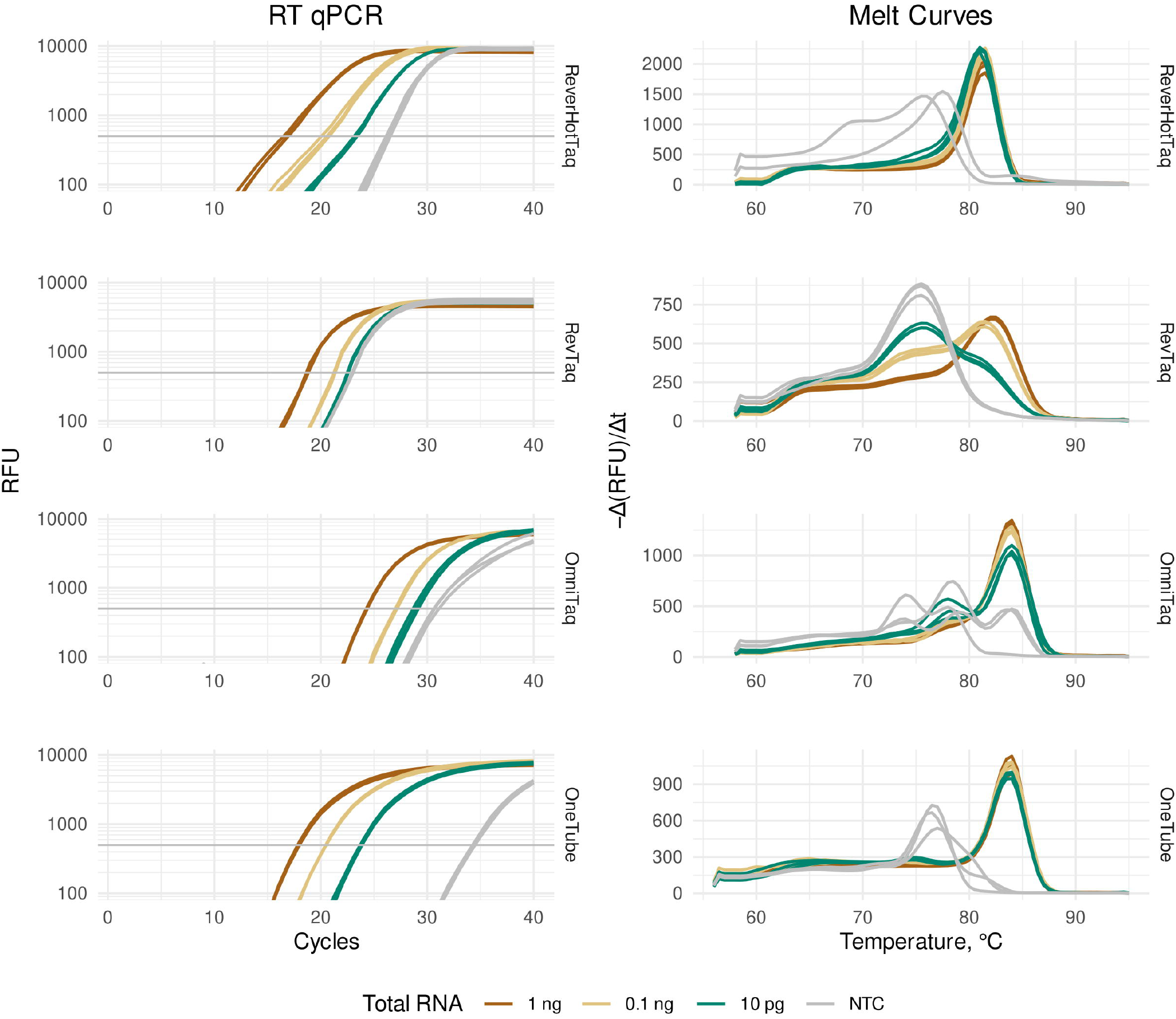
Real-time RT-qPCR with EvaGreen intercalating dye and melting curves of amplicons. The RT-qPCR assays of human *GAPDH* mRNA in the total RNA were carried out with the indicated enzymes: ReverHotTaq, RevTaq, and OmniTaq2 DNA polymerases and OneTube RT-PCRMix. The reaction mixures contained 1 ng (brown curves), 0.1 ng (beige curves), or 10 pg (green curves) of human total RNA per reaction, or no template NTC (gray curves). Melting curves demonstrate the yields of specific (melting peaks over 80 °C) and non-specific (melting peaks below 80 °C) amplicons.

### 3.3 Real-time RT-qPCR with a TaqMan probe

RevTaq, OmniTaq2, and ReverHotTaq DNA polymerases were compared with OneTube RT-PCRMix by performing coupled RT-qPCR assays with a hydrolysis TaqMan probe. The target sequence of SARS-CoV-2 RNA was detected in the sample of nasopharyngeal swab RNA containing SARS-CoV-2 viral gRNA. The reactions were carried out with the series of dilutions of the RNA template, as described in *Materials and Methods*. The results of the assays demonstrated that RevTaq, ReverHotTaq, and OneTube RT-PCRMix had similar efficiency in RT-qPCR with the TaqMan probe (Table 1 and Figure 3). OmniTaq2 DNA polymerase did not generate a fluorescent signal with the TaqMan probe and failed in this test (Fig. 3D).

**Figure 3.**
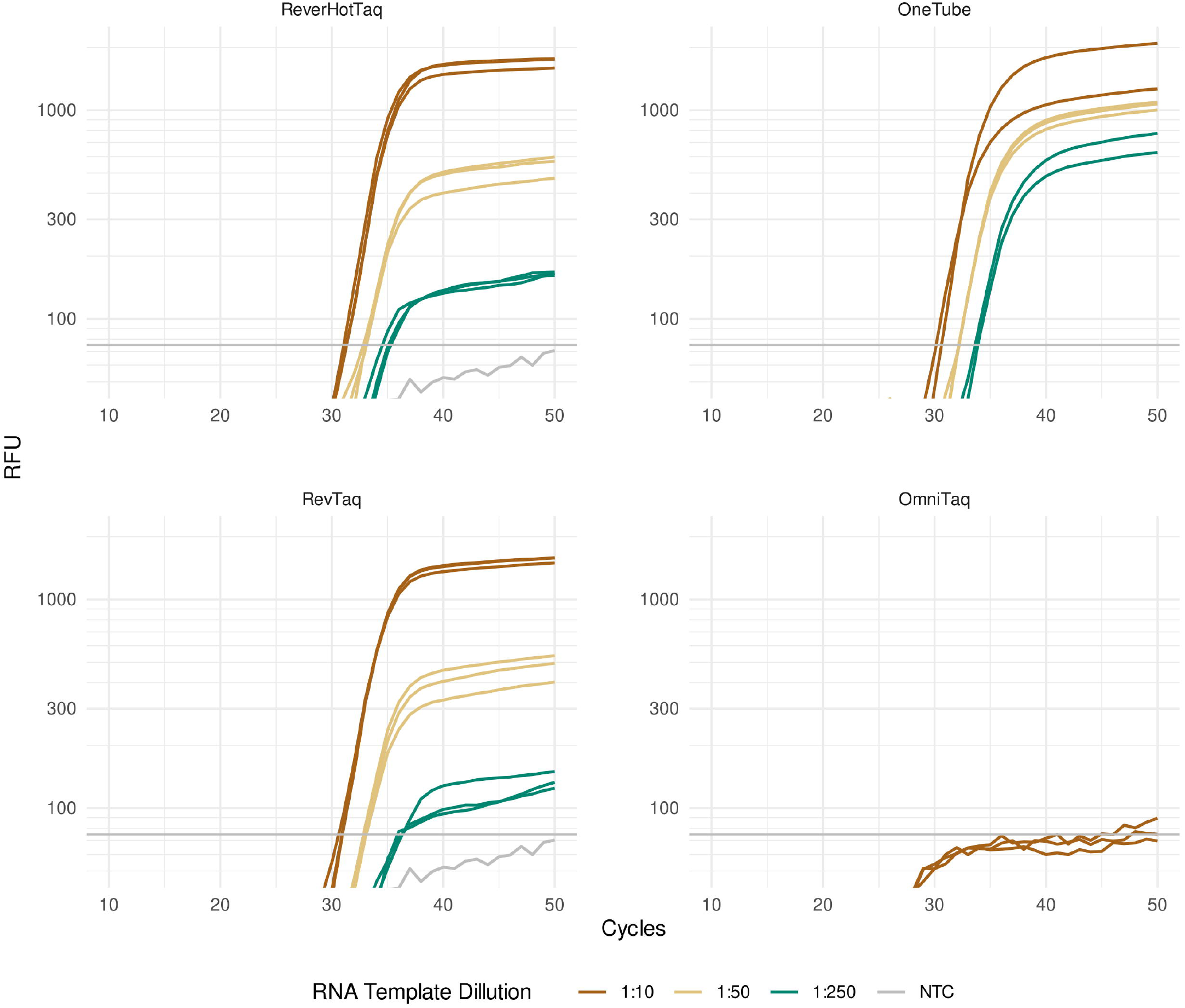
Real-time RT-qPCR with TaqMan probe. The RT-qPCR assays of SARS-CoV-2 viral RNA were carried out with the indicated enzymes: ReverHotTaq, RevTaq, and OmniTaq2 DNA polymerases and OneTube RT-PCRMix. The reaction mixtures contained 10 times diluted (brown curves), 50 times diluted (beige curves), or 250 times diluted (green curves) sample of RNA isolated from SARS-CoV-2-positive nasopharyngeal swabs as a template or no template NTC (gray curves).

## 4 Discussion

RT-PCR and RT-qPCR are widely used methods for the detection of target RNA sequences, including viral RNAs and endogenous mRNAs. In contrast to traditional enzymatic mixtures for RT-PCR that contain a reverse transcriptase and a thermostable DNA polymerase, the modified thermostable RevTaq, OmniTaq2, and ReverHotTaq DNA polymerases with reverse-transcriptase activity could be used as single enzymes for RT-PCR. These artificial polymerases were created by different modifications of Taq DNA polymerase, and therefore, contain different amino acid substitutions. We compared the efficiency of the modified DNA polymerases to the conventional enzyme mixtures for RT-PCR in end-point RT-PCR and real-time RT-qPCR using *B2M* and *GAPDH* mRNAs as the target RNA sequences. These two genes are commonly used for expression studies (Balaji, S. et al., 2020). SARS-CoV2 virus RNA, which was an actual specimen of the viral pathogen, was used as the target RNA template. We used primers and probes that were designed for in-house routine RT-PCR experiments and optimized for applications with conventional RT-PCR enzyme mixtures.

In the end-point RT-PCR experiments, the modified DNA polymerases were compared with the specially designed enzyme mixture for end-point RT-PCR—OneTaq One-Step Enzyme Mix (NEB). We amplified different fragments of the *B2M* mRNA sequence and found that the increasing length of the amplified sequence from 105 to 317 bases decreased the sensitivity of the assays with the modified DNA polymerases in contrast to the standard One-Step Enzyme Mix. This might indicate that the traditional reverse transcriptase from the RT-PCR Kit has better performance with respect to the long RNA fragments than the currently available, modified DNA polymerases. Further, the amplifications with RevTaq polymerase generated visible amounts of non-specific products, in contrast to the other enzymes. The appearance of non-specific products could be the result of the relatively low annealing temperature of the used primers (60 °C). Usually, a high melting temperature is recommended for the primers (>65 °C) used with RevTaq polymerase.

In the real-time RT-qPCR experiments, we compared the modified DNA polymerases with the OneTube RT-PCR Enzyme Mixture specifically designed for coupled RT-qPCR with intercalating dyes or TaqMan probes. When we used EvaGreen dye in the real-time assays, the OneTube RT-PCR Enzyme Mixture, the RevTaq, and the ReverHotTaq polymerases showed better performance than the OmniTaq2. However, the RevTaq polymerase tended to generate non-specific products at low template concentration that prevented target sequence detection. As mentioned above, this could be the result of the relatively low annealing temperature of the used primers that were originally optimized for the conventional OneTube RT-PCRMix containing a reverse transcriptase.

Real-time RT-qPCR with a hydrolysis TaqMan probe is the main technique for the molecular detection of RNA pathogens, including the SARS-CoV-2 virus. We demonstrated that in SARS-CoV-2 RT-qPCR assays with the TaqMan probe, the RevTaq and ReverHotTaq polymerases could be used for the detection of the viral RNA on par with the conventional OneTube RT-PCR Enzyme Mixture. Moreover, RevTaq has been successfully used for SARS-CoV-2 RNA detection in raw patient samples (Kuiper et al., 2020) and wastewater (Babler et al., 2023). The OmniTaq2 DNA polymerase did not generate any fluorescent signal with the TaqMan probe. It could be possible that the OmniTaq2 had a reduced 5’→3’ exonuclease activity in our experiments and could not efficiently hydrolyze the TaqMan probe.

In conclusion, the artificial thermostable DNA polymerases with reverse-transcriptase activity can be used as simple and relative inexpensive tools for the detection of target RNAs by coupled RT-PCR and RT-qPCR. The main disadvantage of the tested polymerases is the low efficiency of long-fragment RT-PCR amplification in comparison to the traditional enzyme mixtures for RT-PCR containing a reverse transcriptase. These artificial polymerases are better suited for the efficient amplification of short (up to 200 bases) RNA sequences, which would be sufficient for a wide spectrum of RT-PCR assays. The RevTaq, OmniTaq2, and ReverHotTaq polymerases could be used for end-point RT-PCR and real-time RT-qPCR with an intercalating dye. Real-time RT-qPCR with a hydrolysis TaqMan probe could be carried out with the RevTaq and ReverHotTaq polymerases.

## 6 Author Contributions

VMK and KBI were responsible for obtaining funding for the research. VMK, and KBI contributed to the conception and design of the study and data curation. DAV, KAB, EVB, EVS, and KBI were involved in the investigation and methodology development. KAB, EVB, EVS, and KBI carried out the formal analysis and validated the results. KBI supervised the research and wrote the first draft of the manuscript. EVS and KBI contributed to manuscript review and revision. KBI was responsible for obtaining necessary resources and preparing visuals.

## 8 Acknowledgments

We thank Dr. Tatiana V. Kramarova from Stockholm University (Stockholm, Sweden), who helped us in the preparation of the article. We would also like to thank the Charlesworth Group (www.cwauthors.com) for English language editing.

## 10 Data Availability Statement

The original contributions presented in the study are included in the article; further inquiries can be directed to the corresponding author.

